# Disruption of Smarce1, a component of the SWI/SNF chromatin remodeling complex, decreases nucleosome stability in mouse embryonic stem cells and impairs differentiation

**DOI:** 10.1101/2022.05.18.492397

**Authors:** Katsunobu Kashiwagi, Junko Yoshida, Hiroshi Kimura, Kyoji Horie

## Abstract

The SWI/SNF chromatin remodeling complex consists of more than 10 component proteins that form a large protein complex of > 1 MDa. The catalytic proteins Smarca4 or Smarca2 work in concert with the component proteins to form a chromatin platform suitable for transcriptional regulation. However, the mechanism by which each component protein works synergistically with the catalytic proteins remains largely unknown. Here, we report on the function of Smarce1, a component of the SWI/SNF complex, through the phenotypic analysis of homozygous mutant embryonic stem (ES) cells. Disruption of Smarce1 induced the dissociation of other complex components from the SWI/SNF complex. Histone binding to DNA was loosened in homozygous mutant ES cells, indicating that disruption of Smarce1 decreased nucleosome stability. Sucrose gradient sedimentation analysis suggested an ectopic genomic distribution of the SWI/SNF complex, accounting for the misregulation of chromatin conformations. Unstable nucleosomes remained during ES cell differentiation, impairing the heterochromatin formation that is characteristic of the differentiation process. These results suggest that Smarce1 guides the SWI/SNF complex to the appropriate genomic regions to generate chromatin structures adequate for transcriptional regulation.

## Introduction

Eukaryotic DNA wraps around histone octamers, each of which contains two copies of Histone H2A, H2B, H3, and H4, to form the nucleosome, a functional unit of chromatin structure [1]. Chromatin is composed of chains of nucleosomes and is packed at various densities related to the transcriptional activity in each region [2]. Active chromatin regions are loosely packed in the nucleus, whereas repressed chromatin regions are tightly packed [3-5]. The formation of chromatin is a benefit for functional storage of nuclear DNA, but it also interferes with the binding of transcription factors to DNA [1, 2]. To overcome this physical interference, eukaryotic cells utilize the energy of ATP hydrolysis to move histones to make chromatin structures suitable for transcriptional regulation [6-8]. These functional changes in chromatin structure are called chromatin remodeling, and these processes are mediated by ATP-dependent chromatin remodeling factor complexes. These complexes have subunits containing a conserved catalytic ATPase domain and are divided into four subfamilies: imitation switch (ISWI), switch/sucrose non-fermentable (SWI/SNF), chromatin helicase DNA binding (CHD), and INO80 or SWR1. All these remodeling complexes commonly change the positions of nucleosomes, but each chromatin remodeling complex also has characteristic functions. ISWI and CHD chromatin remodeling complexes assemble histone octamers and form evenly spaced nucleosomes [9-12]. INO80 subfamily remodelers replace histone H2A-H2B dimer with H2A.Z-H2B dimer [13]. SWI/SNF slides or evicts histones to make a suitable platform for transcriptional regulation [14]. Brm, a catalytic ATPase domain-containing protein of Drosophila SWI/SNF, was originally discovered as a suppressor of Polycomb group protein. Therefore, SWI/SNF is recognized in a broad sense as a Trithorax protein [15].

SWI/SNF chromatin remodeling complexes are composed of more than 10 subunits that form large, species-specific complexes of >1 MDa [16]. Mammalian SWI/SNF chromatin remodeling complexes are related to yeast SWI/SNF and RSC chromatin remodeling complexes in terms of subunit composition. Mammalian SWI/SNF chromatin remodeling complexes are also called BAF, Brg1/Brahma-associated factor complexes [17, 18]. Distinct subfamilies of BAF complexes have been reported in mouse cells and are required to maintain the pluripotent state of undifferentiated cells and their proper differentiation [19-22]. The components of the mouse BAF complexes change during differentiation. The ES cell-specific BAF complex (esBAF) is mainly composed of Smarca4 (Brg1), Arid1a, Smarcb1, a homo-dimer of Smarcc1, Smarcd1/2, Smarce1, Phf10/Dpf2, and actin-like protein 6a [19, 20]. Differentiation of ES cells into post-mitotic neurons accompanies the replacement of the components of esBAF complex: Arid1a by a hetero-dimer of Arid1a and Arid1b, the homo-dimer of Smarcc1 by a hetero-dimer of Smarcc1 and Smarcc2, Phf10/Dpf2 by Dpf1/Dpf3, and Smarcd1/2 by Smarcd1/3. The mutually exclusive catalytic subunits, Smarca4 and Smarca2, are also exchanged during differentiation. The post-mitotic neuron-specific BAF complex is called neuronal BAF (nBAF) [23-26]. BAF complexes are recognized as both a tumor suppressor and oncogene and are the most frequently (∼20%) mutated chromatin regulatory proteins in human cancers [27]. Somatic mutations in human SMARCB1 have been identified in rhabdoid tumor, and loss of SMARCB1 from the canonical BAF complex results in the formation of rhabdoid tumor-specific BAF complex [28]. SMARCE1, thought to be a core component of the BAF complex and is present in all known canonical subfamilies of the BAF complex, is also mutated in meningioma [29-31], and genetic mutation of SMARCE1 causes Coffin–Siris syndrome [32], a multiple congenital anomaly syndrome. A previous study in Drosophila showed that heterozygosity of BAP111, an ortholog of mammalian Smarce1, enhanced the phenotype resulting from partial loss of Brm, a Drosophila homolog of mammalian Smarca2. This indicated that there is a genetic interaction between BAP111 and Brm [33]. Mouse Smarce1 has an HMG domain in its N-terminal domain, which is predicted to direct the BAF complex to bind to appropriate genomic regions [18]. However, it is largely unknown how Smarce1 affects the localization of the BAF complex within the genome, the integrity of the BAF complex, maintenance of a pluripotent state, or differentiation of ES cells.

In the present study, we conducted biochemical and cell biological analyses of Smarce1 using homozygous mutant mouse ES cells. We previously developed a method to rapidly generate homozygous mutant mouse ES cell lines and constructed a homozygous mutant ES cell bank consisting of about 200 mutant cell lines [34]. During the phenotypic screening of the homozygous mutant ES cells, we noticed that mutant ES cells of Smarce1, a component of the BAF complex, exhibit abnormal morphology. We observed an ectopic genomic distribution of mutant cell-specific BAF complex and the induction of instability in nucleosomes. Mutant cells were also impaired in proliferation and showed abnormal differentiation, accompanied by a deficit of heterochromatinization. These results suggest that Smarce1 is required to maintain the integrity the BAF complex and guides the BAF complex to the appropriate genomic regions to form a proper chromatin structure for transcriptional regulation.

## Results

### Smarce1 knockout locally induces H3K9-acetylation in mouse ES cells

The structures of the *Smarce1* alleles of wild-type (*WT*), homozygous mutant (*Smarce1*^*m/m*^), and revertant (*Smarce1*^*r/r*^) ES cells used in this study are shown in Figure 1A. *Smarce1*^*r/r*^ ES cells were obtained by removing the *FRT*-flanked gene trap unit using Flp recombinase as reported previously [34] and were used as a control for the Smarce1 knockout phenotype. Disruption and reversion of Smarce1 were confirmed by Western blot analysis (Fig. 1B).

**Figure 1.**
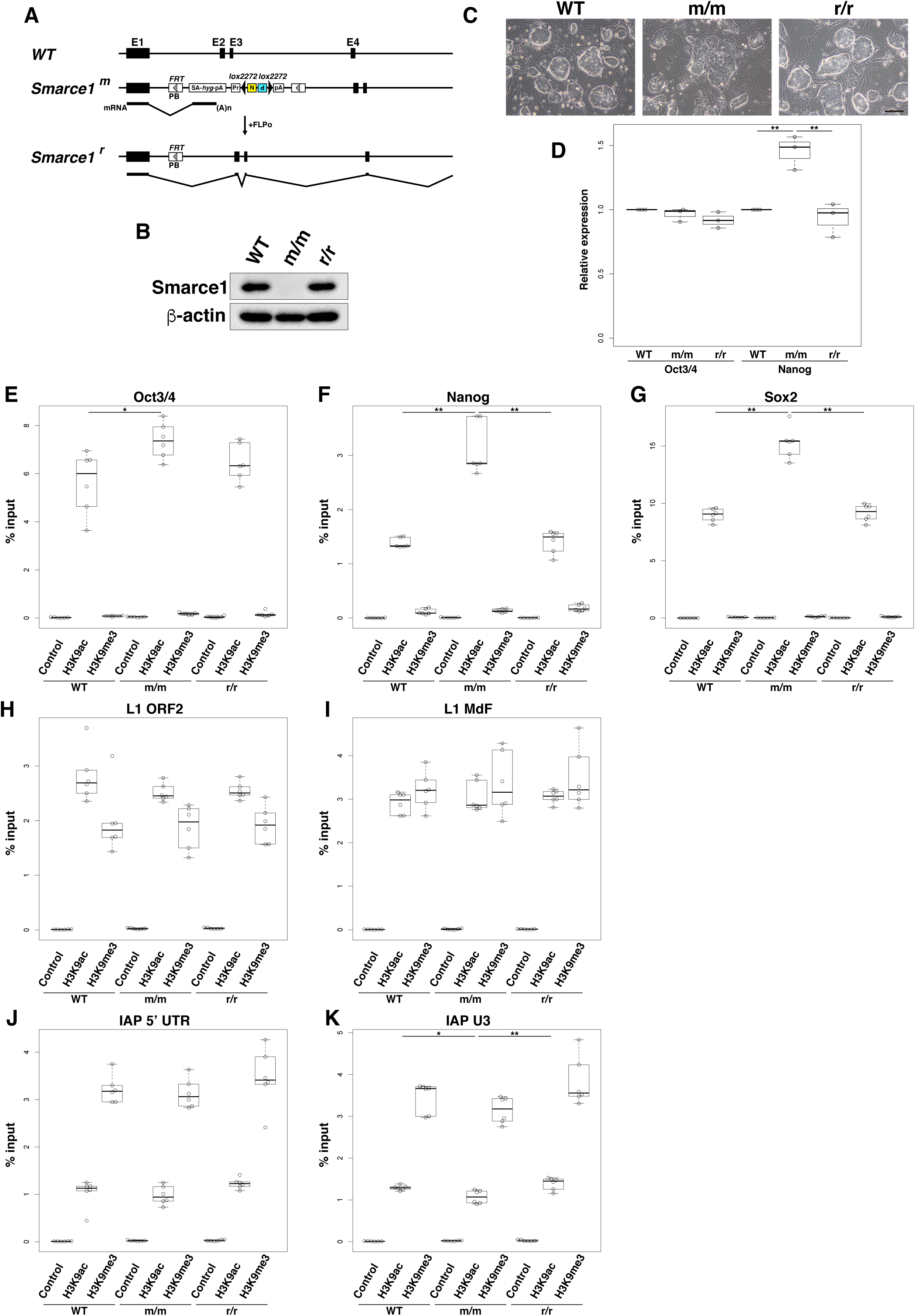
Smarce1 knockout locally induces H3K9-acetylation in mouse ES cells. (A) Structure of the *Smarce1* alleles of wild-type (*WT*), homozygous mutant (*Smarce1*^*m/m*^), and revertant (*Smarce1*^*r/r*^) ES cells used in this study. E, exon; PB, *PiggyBac* transposon; SA, splice acceptor; *hyg*, hygromycin-resistance gene; pA, polyadenylation signal; Pr, *Pgk1* promoter; N, neomycin-resistance gene; P, puromycin-resistance gene. (B) Protein expression analysis of Smarce1. Equal amounts of total proteins (15 μg) from *WT, Smarce1*^*m/m*^, and *Smarce1*^*r/r*^ ES cells were analyzed by immunoblot analysis using the indicated antibodies. *WT*, wild-type; m/m, *Smarce1*^*m/m*^; r/r, *Smarce1*^*r/r*^. (C) Brightfield images of *WT*, homozygous mutant *Smarce1*^*m/m*^, and revertant *Smarce1*^*r/r*^ ES cells. Scale bar, 200 μm. (D) mRNA expression of pluripotency genes in *Smarce1*^*m/m*^ and *Smarce1*^*r/r*^ ES cells relative to *WT* ES cells. Expression levels of *Oct3/4* and *Nanog* were quantified by quantitative RT-PCR and normalized to *β-actin* expression level. Expression levels of *WT* ES cells were set to 1. ** indicates *p-*values of < 0.01. (E)–(K) Chromatin immunoprecipitation assay. Nuclear extracts prepared from *WT, Smarce1*^*m/m*^, and *Smarce1*^*r/r*^ ES cells were incubated with control mouse IgG, anti-H3K9ac, and anti-H3K9me3 antibodies. Immunoprecipitated DNA from specific genomic regions of *Oct3/4* (E), *Nanog* (F), *Sox2* (G), *L1 ORF2* (H), *L1MdF* (I), *IAP 5’ UTR* (J), and *IAP U3* (K) were analyzed by real-time PCR and expressed as a percentage of input DNA. * and ** indicate *p-*values of < 0.05 and < 0.01, respectively.

*WT* ES cells formed round, dome-shaped colonies (Fig. 1C), which is a characteristic feature of undifferentiated mouse ES cells. In contrast, *Smarce1*^*m/m*^ ES cells exhibited flat, irregular shaped colonies (Fig. 1C). *Smarce1*^*r/r*^ ES cells formed round, dome-shaped colonies similar to *WT* ES cells (Fig. 1C), indicating that excision of the gene trap unit reverted the ES cell phenotype. We examined the expression level of pluripotency genes *Oct3/4* and *Nanog*. Although the morphology of *Smarce1*^*m/m*^ ES cells was different from typical undifferentiated ES cells, expression of *Oct3/4* was maintained, and expression of *Nanog* was slightly increased (Fig. 1D). This observation may indicate that the chromatin structure at the pluripotency gene locus is more open in *Smarce1*^*m/m*^ ES cells compared to *WT* cells. To address this possibility, we analyzed the histone modification status of the transcriptional regulatory regions of *Oct3/4, Nanog*, and *Sox2* by chromatin immunoprecipitation followed by real-time PCR (ChIP-qPCR) (Fig. 1E–G). As expected, acetylation of lysine 9 on histone H3 (H3K9ac), a marker for open chromatin [35], was increased in the transcriptional regulatory regions of *Oct3/4, Nanog*, and *Sox2* (Fig. 1E–G). However, there was no significant difference in lysine 9 trimethylation of histone H3 (H3K9me3), which is a marker for heterochromatin [36] (Fig. 1E–G). To investigate whether the alteration of histone modification is a local event or is present genome-wide, we analyzed the retroelements *LINE1* and *IAP* (Fig. 1H–K). *LINE1* and *IAP* are repetitive elements present in the genome at a high copy number and are known to be regulated by histone modifications [37-39]. The levels of H3K9ac and H3K9me3 in the *LINE1* and *IAP* regions were almost the same in *WT, Smarce1*^*m/m*^, and *Smarce1*^*r/r*^ (Fig. 1H–K) except for a slight difference in *IAP U3* (less than 1.3-fold; Fig. 1K), indicating that the alteration of histone modification observed in *Smarce1*^*m/m*^ is present in restricted regions of the genome. Taken together, these data suggest that *Smarce1* knockout induces an open chromatin structure in a local region such as pluripotency genes.

### Smarce1 knockout loosens the binding of histone H3 to DNA

Smarce1 contains an HMG domain, which shares homology with the yeast NHP6A protein [18, 40] (Supplementary Fig. 1). Although yeast NHP6A is not a component of the chromatin remodeling complex, physical and genetic interactions of NHP6A with RSC chromatin remodeling complex have been reported [41]. In addition, NHP6A mutant yeasts have been reported to have loose histone-chromatin binding [42, 43]. These observations suggest the histone-chromatin binding may also be loose in *Smarce1*^*m/m*^ ES cells. To address this possibility, we conducted a biochemical salt extraction assay to examine the binding strength of histones to DNA. Buffers containing different concentrations of salt were added to a nuclear solution of *WT, Smarce1*^*m/m*^, and *Smarce1*^*r/r*^ ES cells to make the final salt concentration 75–450 mM (Fig. 2A). Histone H3 was extracted from these nuclei without cutting the genomic DNA. From the *WT* and *Smarce1*^*r/r*^ nuclei, only a small amount of histone H3 was extracted even at the highest salt concentration (450 mM) (Fig. 2B), indicating a tight association of histone H3 with DNA. In contrast, from *Smarce1*^*m/m*^ nuclei, extraction of histone H3 was increased at moderate salt concentrations (300 mM), and histone H3 was readily extracted at the highest salt concentration (450 mM) (Fig. 2B), indicating a loose association of histone H3 to DNA in *Smarce1*^*m/m*^ nuclei. In accordance with this observation, Arid1a, one of the components of the BAF complex, was also readily extracted from *Smarce1*^*m/m*^ nuclei (Fig. 2B). Extraction of the transcriptional repressor protein Kap1 was also higher in *Smarce1*^*m/m*^ ES cells compared to *WT* and *Smarce1*^*r/r*^ ES cells at the highest salt concentration (450 mM) (Fig. 2B). Unexpectedly, the amount of Kap1 extracted from the nuclei decreased with increasing salt concentration in the extraction buffer (Fig. 2B). Kap1 or a complex containing Kap1 acquired hydrophobicity under high salt concentration and may have been lost from the soluble fraction due to salt precipitation (Fig. 2B). In contrast to these proteins, extraction efficiencies of Smarcc1 and Smarcc2 that were highly and lowly expressed in *WT* ES cells, respectively, did not change between *WT, Smarce1*^*m/m*^ and *Smarce1*^*r/r*^ ES cells (Fig. 2B). These results of the loose association of chromatin proteins with DNA indicate that *Smarce1*^*m/m*^ ES cells have unstable nucleosomes.

**Figure 2.**
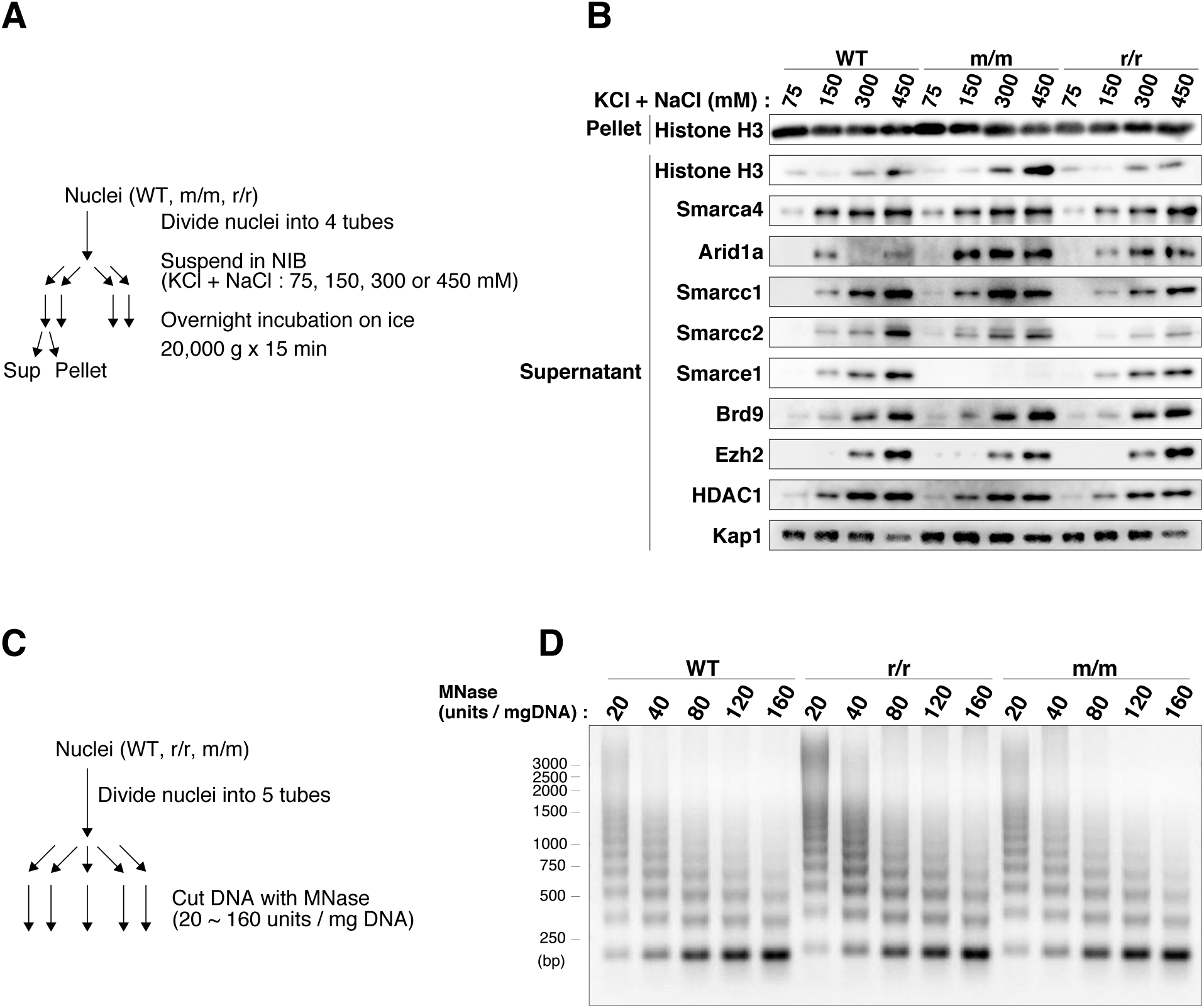
Smarce1 knockout loosens binding of histone H3 to DNA. (A) Schematic representation of the salt extraction assay. Nuclei in a solution containing equal amounts of DNA from *WT, Smarce1*^*m/m*^, and *Smarce1*^*r/r*^ ES cells were treated with buffers of different salt concentrations (75 mM to 450 mM), separated into supernatants and pellets by centrifugation, and analyzed in (B). *WT*, wild-type; m/m, *Smarce1*^*m/m*^; r/r, *Smarce1*^*r/r*^; NIB, nuclei isolation buffer. (B) Association of proteins to chromatin analyzed by immunoblot analysis using the indicated antibodies. Note that histone H3, Arid1a, and Kap1 were more easily extracted in the supernatant fraction. (C) Schematic representation of the MNase sensitivity assay. Nuclei in a solution containing an equal amount of DNA were cut with indicated units of MNase. (D) MNase-treated DNAs were separated on a 1.5% agarose gel and visualized by ethidium bromide staining.

We then analyzed global chromatin architecture by micrococcal nuclease (MNase) sensitivity assay (Fig. 2C). When nuclei isolation and MNase treatment were carried out in the presence of 75 mM salt, in which higher-order chromatin structure is maintained [44, 45], no difference in global digestion pattern of chromatin was observed between *WT, Smarce1*^*m/m*^, and *Smarce1*^*r/r*^ cells (Fig. 2D). This result was consistent with the findings of the salt extraction assay in which histone H3 was tightly associated with chromatin in *Smarce1*^*m/m*^ nuclei in low salt (75 mM) concentration as in *WT* and *Smarce1*^*r/r*^ (Fig. 2B). Taken together, these results indicate that the genome-wide nucleosome positioning is unaffected in *Smarce1*^*m/m*^ ES cells despite weak interactions between histones and DNA.

### The interaction between Smarca4 and the components of the BAF complex is reduced in Smarce1 mutant ES cells

Recent studies have shown that a mutation of *SMARCB* reduced the amount of ARID1A/B and DPF2 in BAF chromatin remodeling complex [46, 47]. These studies suggest that a mutation in one component of BAF chromatin remodeling complex may alter the amount of other components. To explore the possibility that a mutation of *Smarce1* induces changes in the components of esBAF chromatin remodeling complex (Supplementary Fig. 2), we analyzed Smarca4-interacting proteins by immunoprecipitation analysis. Nuclear extracts were prepared from MNase-treated *WT, Smarce1*^*m/m*^, and *Smarce1*^*r/r*^ cells in the presence of 150 mM salt and were immunoprecipitated with anti-Smarca4 antibody at the same salt concentration (Fig. 3). As a control, a normal rabbit IgG was used for a mock immunoprecipitation. Smarca4-interacting proteins were further investigated by immunoblot analysis. Consistent with the results of the salt extraction assay (Fig. 2B), Arid1a and Kap1 were readily extracted from *Smarce1*^*m/m*^ as shown in the input lane (Fig. 3). The amount of Arid1a precipitated with the anti-Smarca4 antibody decreased in *Smarce1*^*m/m*^ (Fig. 3), suggesting a reduction of Arid1a in the BAF complex. Smarca4 successfully pulled down Arid3b, which has not been reported as a component of the BAF complex, even though no protein was detected in the input lane due to limited detection sensitivity. Brd9, a bromodomain-containing protein, has been reported to interact with Smarca4 but not with Smarce1 and to be contained in a non-canonical BAF complex called GBAF complex [26, 47, 48] (Supplementary Fig. 2). Therefore, we investigated the interaction between Smarca4 and Brd9 in *WT, Smarce1*^*m/m*^, and *Smarce1*^*r/r*^ but did not observe any differences between the three cell lines. These results indicate that a Smarce1 deficiency affects the components of the esBAF complex but not the composition of the non-canonical GBAF complex. Smarca4 has also been reported to interact with repressor proteins such as PRC2 protein Ezh2, Kap1 [49] and HDAC1 [50]. Weak interactions of Ezh2 and HDAC1 with Smarca4 were detected in *WT, Smarce1*^*m/m*^, and *Smarce1*^*r/r*^. However, no interaction was detected between Kap1 and Smarca4 in the three cell lines.

**Figure 3.**
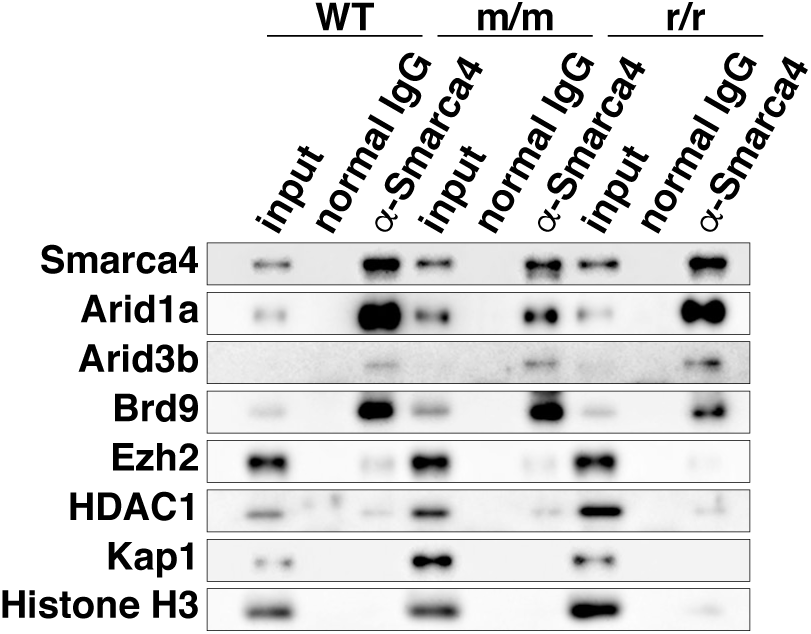
The interaction between Smarca4 and the components of the BAF complex is reduced in *Smarce1*^*m/m*^ ES cells. Immunoblot analysis of Smarca4-associated proteins using the indicated antibodies. Immunoprecipitation was carried out in the presence of 150 mM salt. The input represents 10% of nuclear extracts.

Taken together, the interaction of Smarca4 with Arid1a, a component of the esBAF complex, was decreased in *Smarce1*^*m/m*^ cells. However, the interaction of Smarca4 with components of the GBAF complex, Arid3b, Ezh2, and HDAC1 was unaffected in *Smarce1*^*m/m*^ cells. These results suggest that a deficiency of Smarce1 specifically affects the components of the esBAF complex but not GBAF or the repressor complexes.

### Characterization of the protein composition and genomic distribution of the BAF complex by sucrose gradient sedimentation analysis

To further analyze the properties of the BAF complex in *Smarce1*^*m/m*^ cells, soluble chromatin from MNase-treated *WT* and *Smarce1*^*m/m*^ cells was subjected to 10–40% (W/V) sucrose gradient sedimentation analysis. Fractionated BAF component proteins and other chromatin-associated proteins prepared from *Smarce1*^*m/m*^ cells were compared to those of *WT* cells. We performed experiments with two different salt concentrations: 75 mM and 300 mM. Under 75 mM salt, chromatin is expected to maintain a high-order structure [44, 45]; therefore, interactions between various proteins and chromatin will be detected. Under 300 mM salt, many proteins are expected to dissociate from chromatin.

Under the low salt concentration (75 mM), Smarca4 from *Smarce1*^*m/m*^ cells migrated towards both the top and bottom fractions compared to *WT* cells (Fig. 4A). Other components of the BAF complex, Arid1a, Smarcc1, and Smarcc2, from *Smarce1*^*m/m*^ cells also migrated towards both the top and bottom fractions (Fig. 4A). The molecular weight of Smarce1 is 46.64 kD. Given the distribution of gel filtration molecular markers centrifuged in parallel (Fig. 4A, top), migration of the BAF complex components to the top fractions cannot be explained by the lack of Smarce1 alone. Arid1a protein was detected in fractions 4 and 6 in *Smarce1*^*m/m*^, but not in *WT* (Fig. 4A). From the distribution of the molecular markers, the molecular weight of proteins in fraction 6 would be about 230 kD. Because the molecular weight of Arid1a is 242.05 kD, the Arid1a protein detected in fraction 6 may represent a free protein dissociated from the BAF complex. This observation was consistent with the immunoprecipitation assay (Fig. 3), which suggested that BAF components such as Arid1a dissociated from the complex in *Smarce1*^*m/m*^ ES cells. However, migration to the bottom fractions contradicted the size reduction of the BAF complex. Smarce1 has an HMGB1 domain that has DNA-binding activity [18]. Therefore, when Smarce1 is disrupted, the BAF complex may incorrectly interact with chromatin. Migration of the BAF complex to the bottom fractions suggests the interaction of the BAF complex with heterochromatin regions. This unexpected migration of the BAF complex towards the bottom fractions was accompanied by migration of the PRC2 components Ezh2 and Suz12 to the top fractions (Fig. 4A). Misregulation of Smarca4 localization may have exerted chromatin remodeling ectopically in heterochromatin regions, disrupted the chromatin platform suitable for PRC2 binding, and shifted PRC2 components Ezh2 and Suz12 towards the top fractions. In contrast, Brd9, Arid3b, and other repressor proteins such as HDAC1 and Kap1 did not shift to the bottom or top fractions (Fig. 4A). This observation was consistent with the results of the immunoprecipitation assay showing that the effects of the Smarce1 deficiency were limited to components of the esBAF complex (Fig. 3).

**Figure 4.**
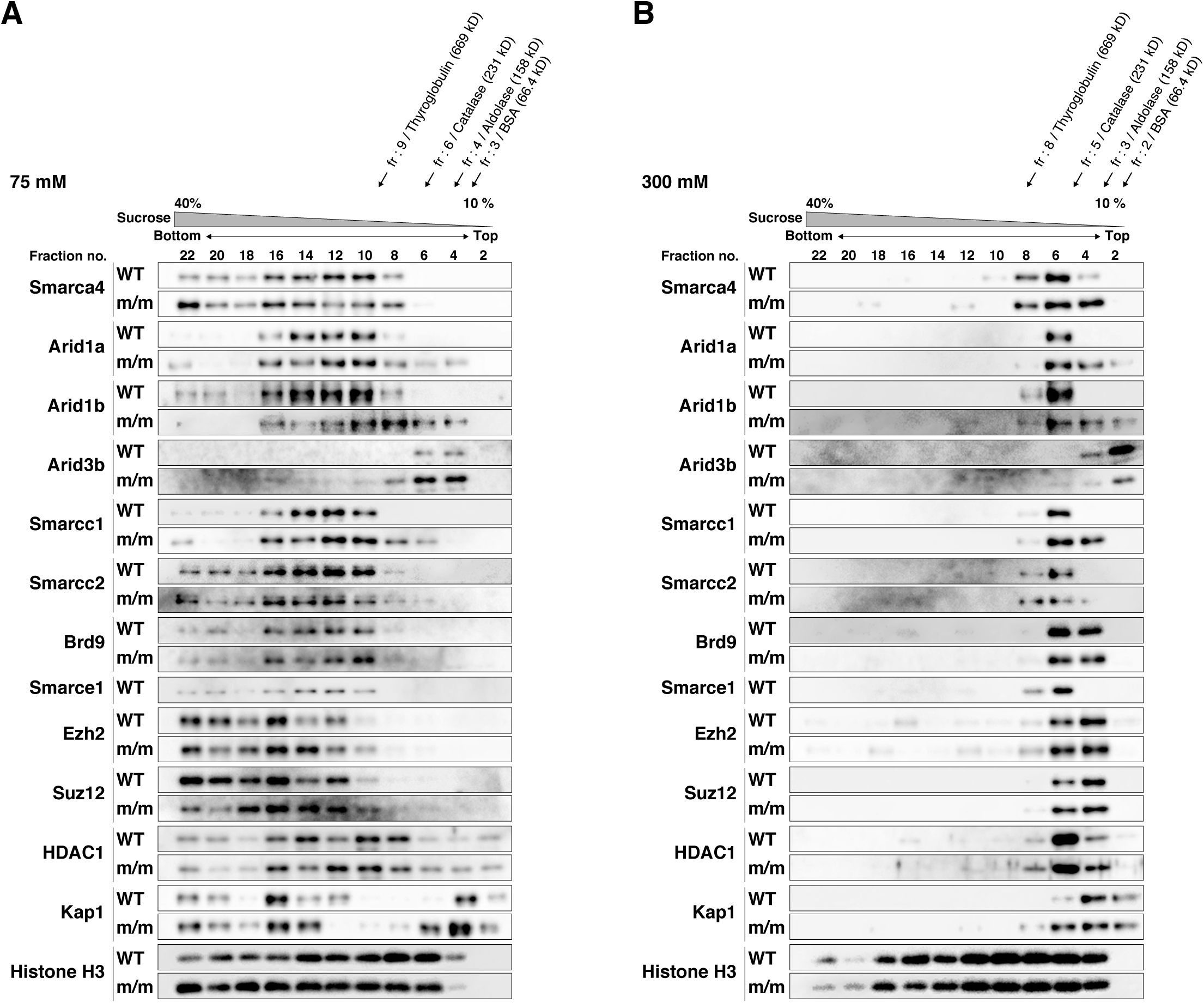
Sucrose gradient sedimentation analysis of the distribution of BAF complex component proteins and chromatin-associated proteins. Nuclear proteins from *WT* or *Smarce1*^*m/m*^ ES cells extracted with 75 mM (A) or 300 mM (B) salt-containing buffer were subjected to 10 to 40% (w/v) sucrose gradient sedimentation analysis. Equal amounts of protein from each fraction were analyzed by immunoblot assay using the indicated antibodies. Estimated molecular weights are shown at the top. *WT*, wild-type; m/m, *Smarce1*^*m/m*^; r/r, *Smarce1*^*r/r*^.

Next, we performed the sucrose gradient sedimentation assay at the high salt concentration (300 mM) for which no interaction between Smarca4 and histone H3 was observed (Supplementary Fig. 3). At this concentration, Smarca4 from *Smarce1*^*m/m*^ cells migrated towards the top fractions, but not to the bottom fractions in contrast to the low salt concentration (Fig. 4B). Other components of the BAF complex, Arid1a, Arid1b, Smarcc1, and Smarcc2 prepared from *Smarce1*^*m/m*^ cells, also co-migrated towards the top fractions only. This observation supports the above-mentioned notion that the migration of the BAF complex components to the bottom fractions under low salt concentration reflects the interaction of the BAF complex with heterochromatin regions. In contrast to esBAF complex component proteins, non-esBAF complex proteins such as Arid3b, Brd9, and repressor proteins were not affected (Fig. 4B), indicating the specificity of the effect of Smarce1-knockout on the esBAF complex.

### Abnormal differentiation of Smarce1 mutant cells is associated with defective heterochromatinization

Undifferentiated ES cells have an open chromatin structure permissive to differentiation stimuli [3]. Upon differentiation stimuli, appropriate genomic regions are heterochromatinized, and a chromatin structure specific to each cell type is established [3, 36]. Abnormal protein composition of the esBAF complex and ectopic distribution of repressor proteins in undifferentiated *Smarce1*^*m/m*^ ES cells suggest that the reorganization of chromatin structure upon differentiation stimuli may be impaired in *Smarce1*^*m/m*^ ES cells. Therefore, we investigated the phenotypes of *Smarce1*^*m/m*^ ES cells during differentiation, with a particular focus on changes in chromatin structure.

*WT, Smarce1*^*m/m*^, and *Smarce1*^*r/r*^ ES cells were cultured in hanging drops for 3 days to form embryoid bodies, and microscopic images were taken for measuring the surface area. Although *WT* and *Smarce1*^*r/r*^ cells developed equally, the surface area of the *Smarce1*^*m/m*^ embryoid bodies was smaller than that of *WT* and *Smarce1*^*r/r*^ (Fig. 5A, B), indicating a delay in the proliferation of mutant cells. To investigate differentiation potential of the *Smarce1*^*m/m*^ cells, embryoid bodies were transferred onto gelatin-coated plates, cultured for an additional 7 days, and stained for mesodermal (α-smooth muscle actin) (Fig. 5C, D) [51] and ectodermal (β-III tubulin) (Fig. 5E, F) [52] markers. *WT* and *Smarce1*^*r/r*^ cells succeeded in differentiating into α-smooth muscle actin-positive cells, and the differentiated cells were square-shaped (Fig. 5D), which is a typical morphology observed in normal differentiation, and appeared throughout the colonies. In contrast, α-smooth muscle actin-positive cells were observed in the peripheral area of the *Smarce1*^*m/m*^ colonies, and they were elongated and rectangular in shape (Fig. 5D). Consistent with the impaired differentiation, more Nanog-positive undifferentiated cells were observed at the center of the *Smarce1*^*m/m*^ colonies than of *WT* and *Smarce1*^*r/r*^ (Fig. 5C). These results indicate the defective differentiation of *Smarce1*^*m/m*^ into mesodermal lineages and the persistence of undifferentiated cells. Regarding the differentiation into ectodermal lineages, β-III tubulin-positive cells were observed at the periphery of the colonies in *WT* and *Smarce1*^*r/r*^, whereas β-III tubulin-positive cells were found within almost the entire region of the colonies in *Smarce1*^*m/m*^ (Fig. 5E, F). Characteristically, neurite-like structures were prominent in *Smarce1*^*m/m*^ colonies and surrounded Nanog-positive cells (Fig. 5E, F), which was not observed in *WT* and *Smarce1*^*r/r*^ cells. A recent study showed enhanced neuronal differentiation in human ARID1A mutant ES cells [53]. The abundant neurite-like structures observed in *Smarce1*^*m/m*^ cells may have been caused by a reduced amount of Arid1a in BAF complex (Fig. 3, 4). Taken together, these results indicate that differentiation into α-smooth muscle actin-positive cells is impaired in *Smarce1*^*m/m*^ cells, but the outgrowth of neurites is enhanced in *Smarce1*^*m/m*^ cells.

**Figure 5.**
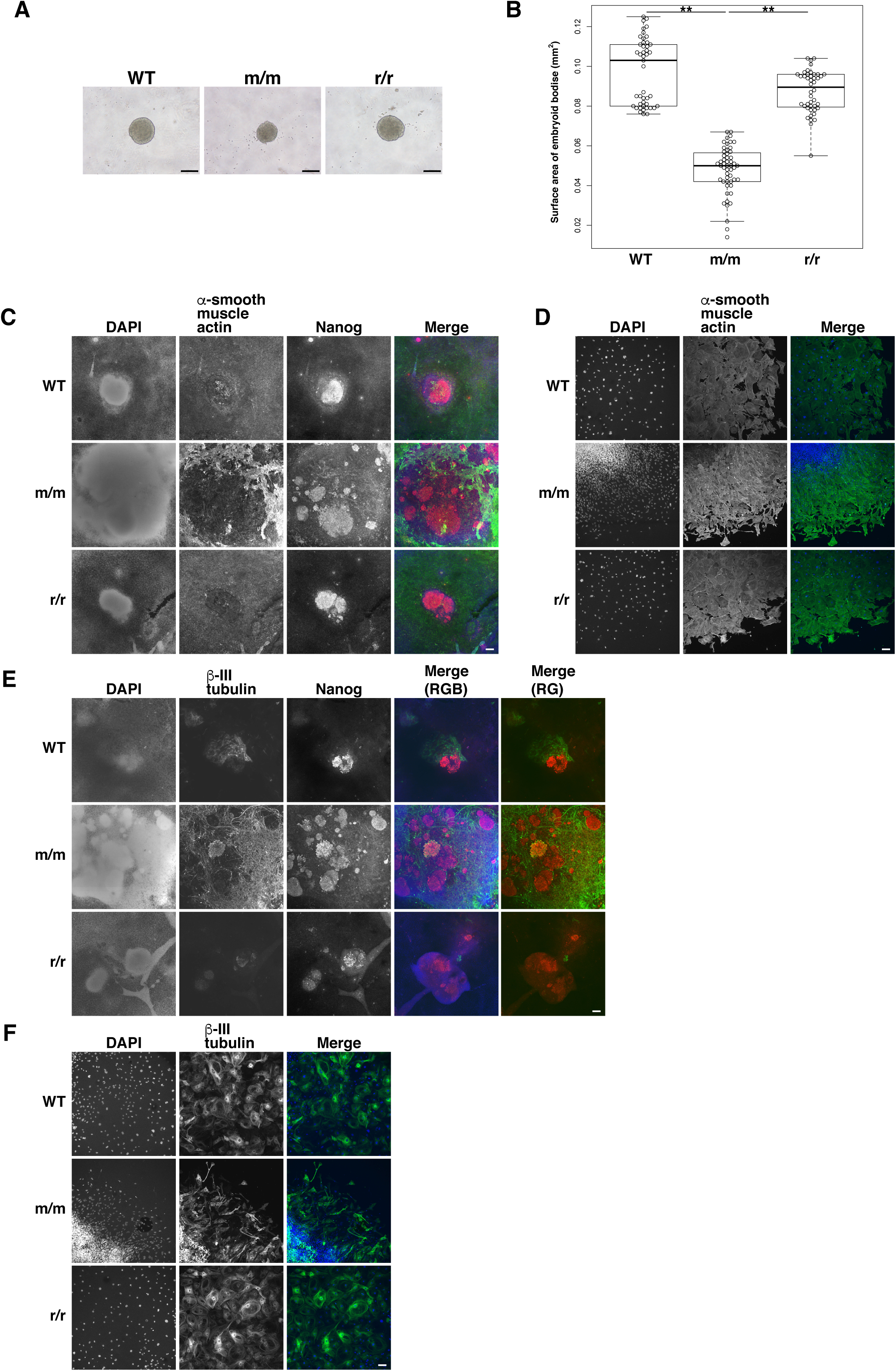
Abnormal differentiation of *Smarce1*^*m/m*^ ES cells. (A)Brightfield images of embryoid bodies of *WT, Smarce1*^*m/m*^, and *Smarce1*^*r/r*^ cells obtained by hanging drop culture (day 3). Images were taken immediately after transferring embryoid bodies onto gelatin-coated coverslips. *WT*, wild-type; m/m, *Smarce1*^*m/m*^; r/r, *Smarce1*^*r/r*^. Scale bars, 200 μm. (B)Quantification of the surface area of embryoid bodies (day 3). ** indicates *p-*values of < 0.01. (C)Immunostaining of differentiated cells with anti-α-smooth muscle actin antibodies on day 10 of differentiation. Embryoid bodies on day 3 (shown in (A)) were cultured on gelatin-coated coverslips for 7 days. Red, green, and blue signals in the merged images represent Nanog, α-smooth muscle actin, and DAPI-stained DNA, respectively. Scale bar, 100 μm. (D)Immunostaining images of α-smooth muscle actin-positive cells on day 10 of differentiation. Green and blue signals in the merged images represent α-smooth muscle actin and DAPI-stained DNA, respectively. Scale bar, 100 μm. (E)Immunostaining images of β-III-tubulin-positive cells around Nanog-positive cells on day 10 of differentiation. Red (R), green (G), and blue (B) signals in the merged images represent Nanog, β-III-tubulin, and DAPI-stained DNA, respectively. Scale bar, 100 μm. (F)Immunostaining images of β-III-tubulin-positive cells in peripheral regions of embryoid bodies on day 10 of differentiation. Green and blue signals in the merged images represent β-III-tubulin and DAPI-stained DNA, respectively. Scale bar, 100 μm.

Next, we investigated heterochromatin formation during ES cell differentiation. Upon the stimulation of differentiation, centromeric heterochromatin foci identified by DAPI-staining increase in number, become smaller, and form discrete structures [54]. These foci show constitutive heterochromatin as evidenced by immunostaining with H3K9me3 [36] and H4K20me3 [55] (Fig. 6A, B). We compared the morphology of these foci in *WT, Smarce1*^*m/m*^, and *Smarce1*^*r/r*^ cells. *WT* and *Smarce1*^*r/r*^ cells formed discrete and round foci (Fig. 6C). In contrast, the foci of *Smarce1*^*m/m*^ cells showed a distorted shape (Fig. 6C). To quantitatively assess the shape of the DAPI-staining foci, we determined the circularity of these foci (Fig. 6D). The circularity of the foci in *Smarce1*^*m/m*^ cells was lower compared to that in *WT* and *Smarce1*^*r/r*^ cells, suggesting the impaired formation of constitutive heterochromatin.

**Figure 6.**
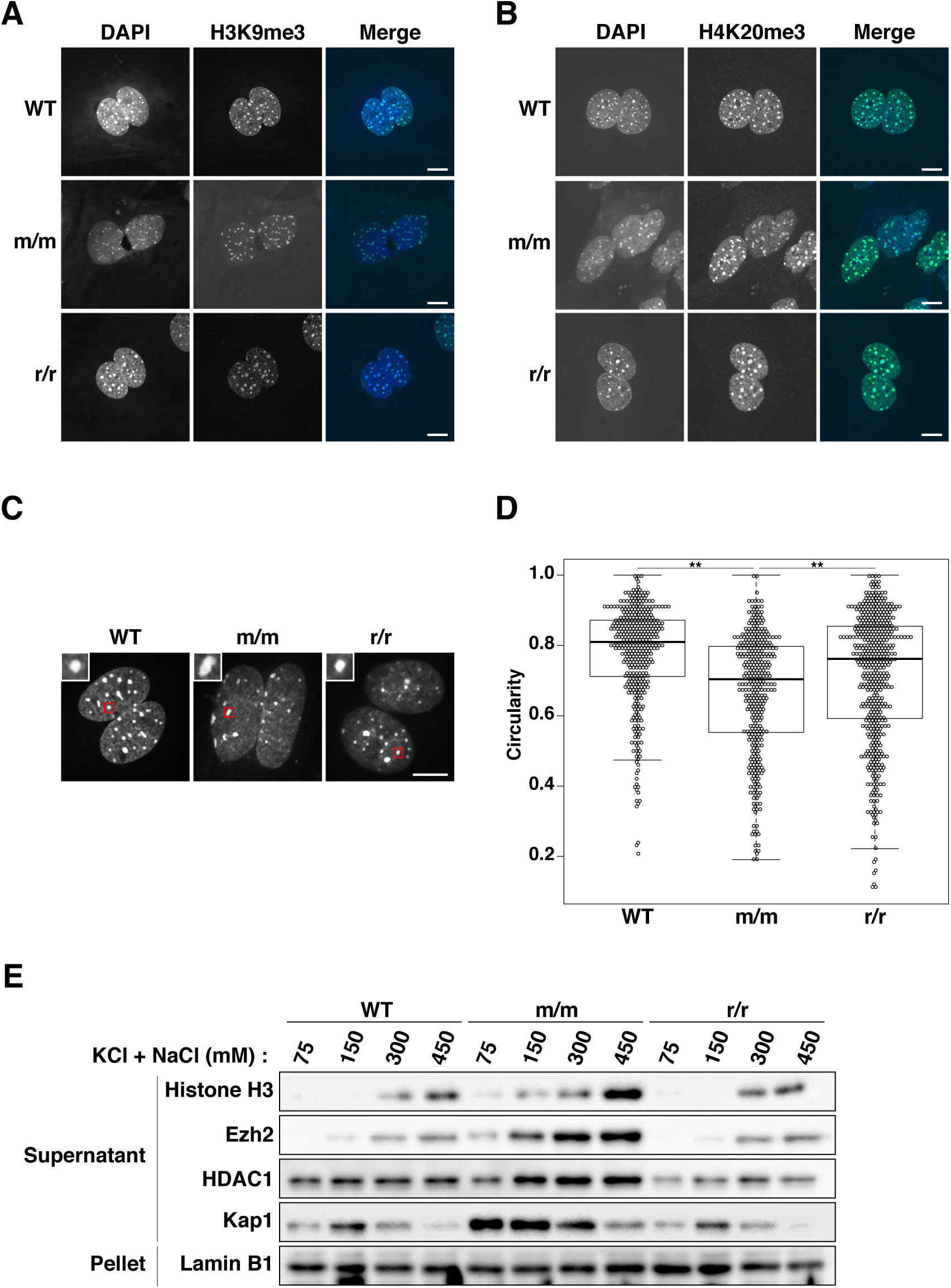
Defects in heterochromatin formation during the differentiation of *Smarce1*^*m/m*^ ES cells. (A, B) Immunostaining of heterochromatin markers on day 10 of differentiation. Green and blue signals in the merged images represent H3K9me3 and DNA in (A) and H4K20me3 and DNA in (B), respectively. Scale bar, 10 μm. (C) Nuclei images of *WT, Smarce1*^*m/m*^, and *Smarce1*^*r/r*^ cells. DAPI foci highlighted by red squares are presented as enlarged images on the top left side. Scale bar, 10 μm. (D) Circularity of DAPI foci in *WT, Smarce1*^*m/m*^, and *Smarce1*^*r/r*^ cells. Circularity was measured by 4π(area/perimeter^2). ** indicates *p-*values of < 0.01. (E) Loosened association of proteins to chromatin by salt extraction assay. Each protein was detected by immunoblot analysis using the indicated antibodies.

To further confirm the impaired formation of heterochromatin in differentiated cells, we conducted a biochemical analysis (Fig. 6E). Histone H3 and the repressor proteins Kap1, Ezh2, and HDAC1 were extracted from the nuclei of differentiated cells at various salt concentrations. A greater quantity of histone H3 was extracted from *Smarce1*^*m/m*^ cells than from *WT* and *Smarce1*^*r/r*^, suggesting loose chromatin structure in *Smarce1*^*m/m*^ cells (Fig. 6E). Consistent with the morphological abnormality of the constitutive heterochromatin foci (Fig. 6A–D), Kap1 was more readily extracted from *Smarce1*^*m/m*^ cells than *WT* and *Smarce1*^*r/r*^ cells (Fig. 6E). As observed in undifferentiated ES cells (Fig. 2B), the amount of Kap1 extracted from the nuclei decreased with increasing salt concentration in the extraction buffer (Fig. 6E). Ezh2, an integral component of PRC2 that regulates facultative heterochromatin, and HDAC1, which is associated with both constitutive and facultative heterochromatins, were also more readily extracted from *Smarce1*^*m/m*^ cells than *WT* and *Smarce1*^*r/r*^ cells (Fig. 6E), suggesting that heterochromatin formation was broadly impaired.

Based on these observations, we speculated that weak binding of histones and repressor proteins to chromatin was responsible for impaired heterochromatin formation in *Smarce1*^*m/m*^ cells (Fig. 7).

**Figure 7.**
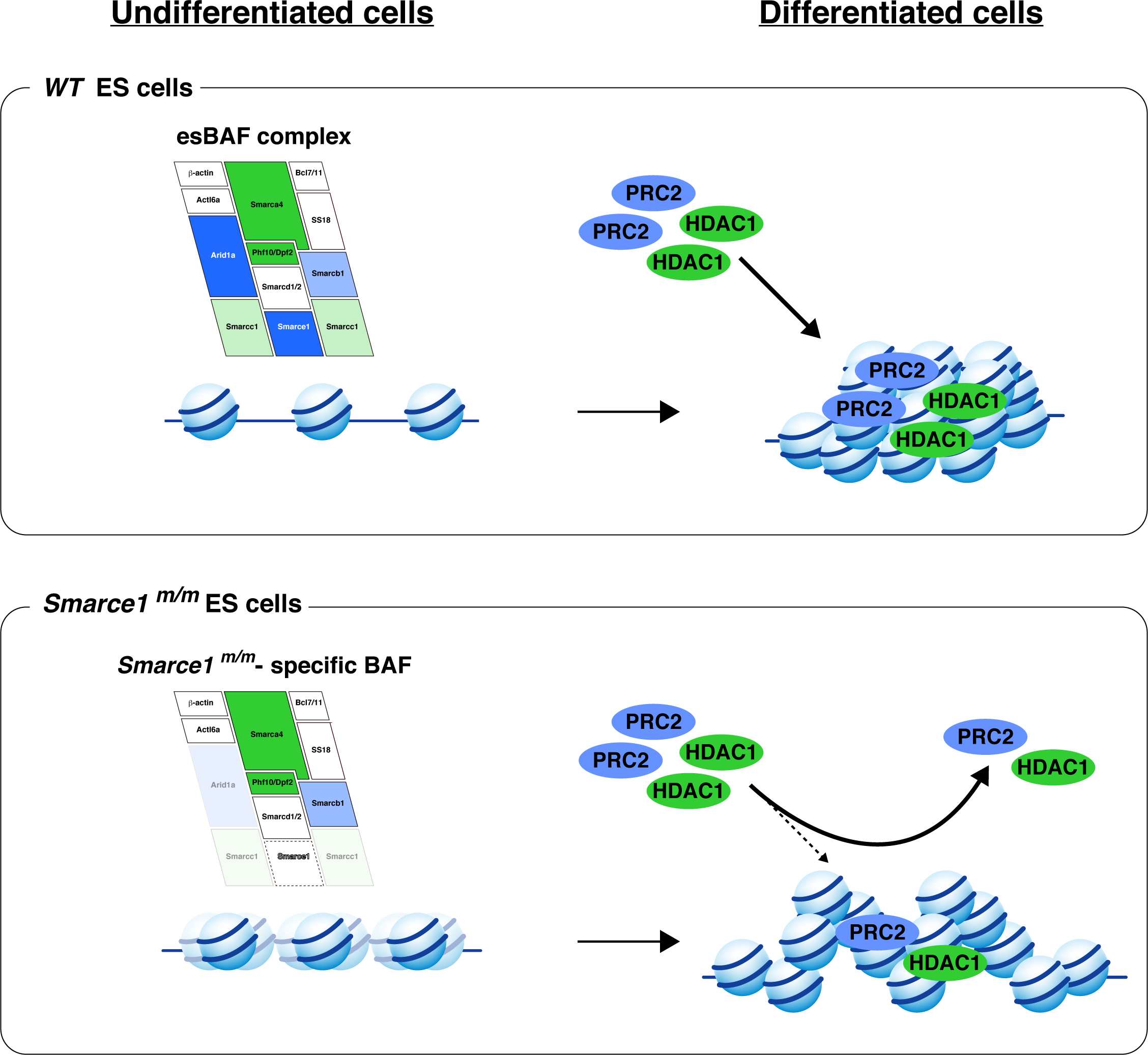
Hypothetical model of the induction of unstable chromatin structure in *Smarce1*^*m/m*^ cells. Histone dissociation from chromatin was induced in undifferentiated *Smarce1*^*m/m*^ ES cells by ectopic genomic localization of the *Smarce1*^*m/m*^-specific BAF complex, leading to unstable chromatin structure. During ES cell differentiation, the unstable chromatin structure might not be a suitable platform for binding of repressor proteins such as PRC2 and HDAC1. Impaired recruitment of repressor proteins to proper genomic regions might further induce abnormal heterochromatinization and differentiation.

## Discussion

The current study showed that disruption of Smarce1 decreases nucleosome stability in mouse ES cells and impairs heterochromatin formation during differentiation (Fig. 7). Smarce1 contains an HMG domain that has been shown to interact with DNA [18]. Other components of the esBAF complex, such as Arid1a, Smarcb1, Smarca4, and Dpf2, also contain DNA-[28, 56-62] or histone binding domains [62-66]. The genomic distribution of the BAF complex is thought to be determined by the overall effect of these BAF complex components. Since the genomic distribution of the BAF complex seemed altered in the absence of Smarce1 as determined by the sucrose gradient sedimentation assay (Fig. 4A), we speculate that Smarce1 serves as a guide for placing the BAF complex in the appropriate genomic regions. We hypothesize that the *Smarce1*^*m/m*^-specific BAF complex may exert remodeling effects on ectopic genomic regions, slide histones along the DNA, and induce the loosening of chromatin structure (Fig. 7). It has been reported that loosely structured chromatin is not suitable for the nucleosome binding of Polycomb group proteins [67, 68]. Recent studies have also shown that ectopic recruitment of the BAF complex to chromatin to which Polycomb group proteins are already bound leads to the release of Polycomb group proteins [69, 70]. Therefore, we speculate that the enhanced release of Polycomb group proteins from chromatin observed in *Smarce1*^*m/m*^ nuclei (Fig. 4A, 6E) was caused by the ectopically distributed *Smarce1*^*m/m*^-specific BAF complex.

Mutation of BAF complex components induces tumorigenesis [16, 26, 27, 71, 72]. For example, SS18, a component of the BAF complex, is reported to fuse to SSX Family Member 2 (SSX) by chromosomal translocation and causes synovial sarcoma. The mutant BAF complex containing this SS18-SSX fusion protein evicts PRC2 from *PAX3* and *SOX2* loci, decreases H3K27me3 levels, and increases the expression of these genes [73]. A mutation of SMARCE1 has been reported to cause meningiomas [29-31]. Given the similarities to synovial sarcoma formation, the ectopic distribution of the BAF complex and the eviction of PRC2 observed in *Smarce1*^*m/m*^ in the current study (Fig. 4A, 6E) may be responsible for meningioma formation.

The HMG domain of mouse Smarce1 shares homology with the HMG box of yeast NHP6A and NHP6B (Supplementary Fig. 1) [18]. NHP6A and NHP6B physically and genetically interact with the yeast RSC chromatin remodeling complex that is closely related to the mammalian BAF complex [17, 18]. In addition, the synthetic lethality of double mutations of the yeast catalytic subunit of the RSC complex and NHP6A/B indicates the genetic interaction between these factors [74]. Furthermore, it has been reported that the association of histone to chromatin was loosened in yeast NHP6A mutant cells [42, 43], which resembled our finding in *Smarce1*^*m/m*^ cells. A yeast ortholog protein of mouse Smarce1 has not been reported thus far. Similarities between yeast NHP6A and mouse Smarce1 suggest that NHP6A may be the functional yeast counterpart of mouse Smarce1.

We observed a reduced association of Arid1a to the BAF complex in *Smarce1*^*m/m*^ (Fig. 3, 4). Interestingly, a reduction of Smarce1 in the BAF complex was reported in Arid1a mutant cells [58]. These observations suggest a strong interaction between Smarce1 and Arid1a. A recent study shows that the prior presence of Smarce1 in the core of the BAF complex is required for the efficient recruitment of Arid1a to form the canonical BAF complex [75]. The impaired association of Arid1a with the BAF complex observed in *Smarce1*^*m/m*^ in the current study supports this concept. As mentioned above, both Smarce1 and Arid1a possess a DNA-binding domain [18, 56-58, 76]. The combined loss of the two DNA-binding domains in *Smarce1*^*m/m*^ may exacerbate the misregulation of the BAF complex and contribute to various phenotypes such as the formation of meningioma in humans.

Accumulation of histone acetylation was detected in the transcriptional regulatory regions of the pluripotent factors Nanog, Oct3/4, and Sox2 (Fig. 1E–G). In contrast to these regions, enrichment of histone acetylation at the loci of the retroelements IAP and LINE1 was minimal, if detected at all, in *Smarce1*^*m/m*^ (Fig. 1H–K) even though the repressor protein Kap1, which has been reported to repress these retroelements [77], readily dissociated from chromatin in the salt extraction assay (Fig. 2B). These results suggest that although Kap1 was easily extracted from *Smarce1*^*m/m*^ nuclei, an additional, unidentified silencing mechanism exists for IAP and LINE1.

After the differentiation of ES cells, morphologically more mature neuronal cells were observed in *Smarce1*^*m/m*^ compared with *WT* and *Smarce1*^*r/r*^ (Fig. 5E, F). As described above, we observed a reduction of Arid1a from the BAF complex of *Smarce1*^*m/m*^ cells (Fig. 3, 4). A previous report showed enhanced neuronal differentiation of ARID1A knockout human ES cells due to an impaired interaction between ARID1A and REST, a repressor of neuronal differentiation [53]. Furthermore, human SMARCE1 has been reported to interact with REST and is required for REST-mediated repression of neuronal genes [78]. Based on these reports, we speculate that the function of Rest was impaired in *Smarce1*^*m/m*^ cells because of the reduction of Arid1a and complete loss of Smarce1 in the BAF complex, thus leading to the enhanced neuronal differentiation (Fig. 5E, F). Both SMARCE1 and ARID1A are causative genes for Coffin-Siris syndrome [32], a multiple congenital anomaly syndrome. The impaired proliferation and abnormal differentiation of *Smarce1*^*m/m*^ ES cells observed in the present study (Fig. 5) may be associated with some of the developmental disorders of Coffin-Siris syndrome.

Our observations in Smarce1 mutant cells revealed not only the role of Smarce1 for maintaining the BAF complex integrity but also the functions of the BAF complex itself in the formation of a suitable chromatin environment for transcriptional regulation in undifferentiated and differentiated cells. Further studies using mutant cells of other components of the BAF complex will help to elucidate new functions of each component in the maintenance of the BAF complex integrity and chromatin structure formation.

## Methods

### Construction of the gene trap vector and insertion site in the Smarce1 gene

A Smarce1-heterozygous mouse ES cell clone (*Smarce1*^*m*^) was obtained using the piggyBac transposon-based gene trap vector containing the same gene trap unit we used previously [34]. The piggyBac gene trap vector was generated as follows. First, a 0.82-kb BglII-ApaI fragment of pT2F2GFP [34] containing the FRT-flanked GFP gene was inserted into the BglII-ApaI site of pPB-MCS-P5 [79], resulting in pPB-F2GFP. Next, a 4.8-kb XhoI-PmlI fragment of the Tol2 gene trap vector pT2F2-SAhygpA-N22 [34] was cloned into the XhoI-PmlI site of the pPB-F2GFP located between the two inverted terminal repeats of the piggyBac transposon, resulting in pPB-SAhygA-NP22. Gene trapping was conducted as described previously [34] and the ES cell clone containing the vector insertion at the first intron of the Smarce1 gene was identified. The flanking genomic sequence of the vector insertion site is 5’-TTAATCGCCCCGAGACTGTTTTCTTCC-3’.

### Cell culture

*Smarce1* homozygous mutant ES cells (*Smarce1*^*m/m*^) were obtained by doxycycline-induced interchromosomal recombination as described previously [34]. Revertant ES cells (*Smarce1*^*r/r*^) were obtained by excising the gene trap unit using Flp-mediated recombination as described previously [34]. Embryoid bodies (EBs) were formed in hanging drops containing 1,000 cells in 20 μl media in the absence of leukemia inhibitory factor (LIF) on the lid of a culture dish and cultured for 3 days. EBs were cultured for a further 7 days on gelatin-treated coverslips. For the salt extraction assay, differentiation was induced by culturing ES cells (4.0 × 10^5^ cells) on a low attachment cell culture dish (Greiner, CELLSTAR, Cell-Repellent Surface, 628979) to form EBs in the absence of LIF for 3 days. For further induction of differentiation, EBs were cultured on a gelatin-treated dish for another 7 days. After transferring EBs to the gelatin-treated coverslips or culture dishes, the differentiation medium was changed every 2 days.

### Preparation of total cell extract

Cells were trypsinized and washed with PBS. The cell pellet was solubilized with 8 M urea containing 0.1 M NaH_2_PO_4_, 10 mM Tris–HCl (pH 8.0), cOmplete™ EDTA-free protease inhibitor cocktail (Roche, 11873580001), and 0.1 mM phenylmethylsulfonyl fluoride (PMSF). The amount of protein was measured by the Bradford method (Bio-rad, 500-0001) using BSA as a standard. An equal amount (15 μg) of each protein was subjected to immunoblot analysis as described below.

### Nuclei preparation

Undifferentiated and differentiated ES cell nuclei were prepared as described elsewhere with some modifications [80]. Cells were washed with PBS and treated with trypsin for dissociation. Trypsin treatment was terminated by adding 10% calf serum-containing medium. Cells were harvested by centrifugation at 300 × g for 5 min at room temperature and washed with PBS. Cells were collected again as described above, resuspended, and washed with ice-cold nuclei isolation buffer (NIB) containing 10 mM Tris-HCl (pH 7.5), 60 mM KCl, 15 mM NaCl, 1.5 mM MgCl_2_, 1 mM CaCl_2_, 0.25 M sucrose, 10% (v/v) glycerol, 10 mM sodium butyrate, 1 mM dithiothreitol (DTT), 0.1 mM PMSF, and cOmplete™ EDTA-free protease inhibitor cocktail. Cells were collected by centrifugation at 300 × g for 5 min at 4ºC and re-suspended in NIB. An equal volume of NIB containing 0.2% (v/v) NP40 buffer was then added to cell suspensions to bring the final concentration of NP40 to 0.1% (v/v). Cells were incubated on ice for 10 min and centrifuged at 300 × g for 5 min at 4ºC. Supernatants containing cytoplasmic proteins were discarded. Pelleted nuclei were resuspended in NIB and centrifuged again at 300 × g for 5 min at 4ºC. Finally, nuclei were resuspended in NIB.

### Salt extraction assay

Nuclei were collected as described above, and a small amount of nuclei solution was taken into saturated 5 M NaCl, 8 M Urea buffer to measure the DNA concentration by UV absorbance at 260 nm (20 OD_260_ units corresponded to 1 mg/ml DNA) [81]. The DNA concentration of the nuclei solution was adjusted to 1.5 mg/ml DNA with NIB. An equal number of nuclei in NIB was divided into four tubes and extracted with an equal volume of nuclei extraction buffer (NEB) containing 10 mM Tris-HCl (pH 7.5), 10 mM EDTA, 0.25 M sucrose, 10% (v/v) glycerol, 10 mM sodium butyrate, 1 mM DTT, 0.1 mM PMSF, cOmplete™ EDTA-free protease inhibitor cocktail, and different concentrations of NaCl (75, 225, 525, or 825 mM NaCl). The resulting salt (KCl with NaCl) concentration of each tube was 75, 150, 300, or 450 mM, respectively. After overnight incubation on ice, nuclei were subjected to centrifugation at 20,000 × g for 15 min at 4ºC. The supernatant fraction was collected, and the nuclear pellet was dissolved in 8M urea buffer containing 0.1 M NaH_2_PO_4_, 10 mM Tris-HCl (pH 8.0), 0.1 mM PMSF, and cOmplete™ EDTA-free protease inhibitor cocktail. Equal samples in terms of initial nuclei number of each fraction were subjected to SDS-PAGE and then analyzed by immunoblot analysis as described below.

### MNase sensitivity assay

An equal number of nuclei in NIB was divided into five tubes and was pre-incubated at 30ºC for 10 min. The nuclei were treated with 20, 40, 80, 120, or 160 units/mg DNA of MNase at 30ºC for 10 min. The reaction was terminated by adding EDTA to a final concentration of 5 mM. The MNase-treated DNA samples were treated with 20 μg/ml RNase at 37ºC for 1 h and then 40 μg/ml proteinase K at 56ºC overnight. On the following day, DNA samples were further extracted twice with 25:24:1 phenol-chloroform-isoamyl alcohol and then extracted once with chloroform-isoamyl alcohol. The extracted DNA samples were precipitated with ethanol and analyzed by 1.5% agarose gel electrophoresis in 1× TAE buffer. The DNA was visualized with ethidium bromide using a UV trans-illuminator.

### Purification of Smarca4-associated proteins

Isolated nuclei in NIB were pre-incubated at 30ºC for 10 min and subjected to MNase (20 units/mg DNA) treatment at 30ºC for 10 min. After MNase treatment, NEB225 or NEB525 containing 225 mM NaCl or 525 mM NaCl was added to the nuclei solution and incubated overnight on ice. The resulting salt (KCl with NaCl) concentration of the nuclei solution was 150 mM or 300 mM, respectively. The nuclear extract was separated by centrifugation at 12,800 × g at 4ºC for 10 min. An equal volume of NEB150 or NEB300 containing 150 mM NaCl or 300 mM NaCl with 0.2% (v/v) NP40 was added to the nuclear extract to bring the final NP40 concentration to 0.1% (IP buffer). Antibodies against mouse Smarca4 (5 μg, Abcam 110641) were added to the nuclear extract and then incubated overnight at 4ºC with rotation. As a negative control, an equal amount of normal rabbit IgG (MBL, PM035) was added to the nuclear extract. The next day, Dynabeads protein G (Thermo Fisher Scientific, 10003D) pre-equilibrated with IP buffer were added to the nuclear extract and incubated at 4ºC for 4 h with rotation. Proteins that did not bind to the Smarca4 antibodies were separated by placing the Smarca4 associated proteins-Dynabeads complexes on a magnet. The complexes were washed three times with IP buffer at 4ºC with rotation for 10 min. Smarca4-associated proteins were collected by placing the complexes on a magnet and eluted with Laemmli SDS-PAGE sample buffer [82]. The eluted samples were subjected to SDS-PAGE and immunoblot analysis as described below.

### Sucrose gradient sedimentation

Isolated nuclei were treated with MNase (20 units/mg DNA) and extracted with NEB75 or NEB525 on ice overnight. The resulting salt concentration in each extract was 75 and 300 mM, respectively. The next day, the extracts were subjected to centrifugation at 12,800 × g for 10 min at 4ºC. The supernatants were further overlaid onto10–40% (w/v) sucrose gradient buffer containing NEB75 or NEB300 and centrifuged at 50,000 rpm for 3 h at 4ºC using a TLS-55 rotor (Beckman). After centrifugation, equal volumes of each fraction were collected from the top of the centrifugation tube. The fractionated samples were mixed with Laemmli SDS-PAGE sample buffer and subjected to SDS-PAGE and immunoblot analysis as described below.

### Immunoblot analysis

Protein samples dissolved in Laemmli buffer were separated on SDS-PAGE gel and transferred to Immobilon-P PVDF membrane (Millipore, IPVH00010). Transferred protein samples were detected using following primary antibodies: anti-Smarca4 (1:2000; Abcam, 110641), anti-Arid1a (1:2000; Cell Signaling Technology, 12354), anti-Arid1b (1:2000; Cell Signaling Technology, 92964), anti-Arid3b (1:2000; Bethyl Laboratory Inc., A302-565A), anti-Smarcc1 (1:2000; Cell Signaling Technology, 11956), anti-Smarcc2 (1:2000; Cell Signaling Technology, 12760), anti-Smarce1 (1:2000; Cell Signaling Technology, 33360), anti-Brd9 (1:2000; Active Motif, 61537), anti-Ezh2 (1:2000; Cell Signaling Technology, 5246), anti-Suz12 (1:2000; Cell Signaling Technology, 3737), anti-HDAC1 (1:2000; Millipore, 06-720), anti-Kap1 (1:5000; Active Motif, 61173), anti-LaminB1 (1:400; Santa Cruz, Sc-20682), anti-β actin (1:4000; Sigma, A5441), and rat anti-Histone H3 serum (1:8000; provided by H. Kimura). Membrane-bound primary antibodies were detected using horse radish peroxidase conjugated anti-rabbit IgG (Cytiva, NA934), anti-mouse IgG (Cytiva, NA931), and anti-rat IgG (Bethyl, A110-305P). Immunoreactive signals were detected using Chemi-Lumi One L (Nacalai Tesque, 07880), Chemi-Lumi One Ultra (Nacalai Tesque, 11644), or ECL prime Western Blotting Detection Reagent (Cytiva, RPN2232).

### Immunofluorescence

EBs were seeded on 0.1% (w/v) gelatin-coated coverslips. Cells were fixed with 4% paraformaldehyde, 100 mM HEPES-HCl (pH 7.4) buffer for 20 min at room temperature and were washed twice with PBS. After fixation, cells were permeabilized with 0.5% (v/v) Triton X-100 in PBS for 20 min at room temperature and were washed with PBS. Cells were further blocked with Blocking One-P (Nacalai Tesque, 05999-84) for 20 min at room temperature and then incubated overnight at 4ºC with the following primary antibodies in antibody dilution buffer (PBS containing 1/10 × Blocking One-P) as indicated: anti-H3K9me3 (1:1,000; 2F3, provided by H. Kimura), anti-H4K20me3 (1:1,000; 27F10, provided by H. Kimura), anti-β-III tubulin (1:125; R&D Systems, MAB1195), anti-α smooth muscle actin (1:250; Sigma-Aldrich, A5228), and anti-Nanog (1:250; ReproCell, RCAB004P-F). After 6 washing steps with PBST (PBS with 0.1% (v/v) Tween-20) for 5 min each, cells were incubated with the following fluorescence-conjugated secondary antibodies in antibody dilution buffer as indicated: Goat anti-mouse IgG Highly cross-adsorbed secondary antibody, Alexa Fluor 488 (1:1,000; Thermo Fisher Scientific, A-11029) and Goat anti-rabbit IgG Highly cross-adsorbed secondary antibody, Alexa Fluor 594 (1:1,000; Thermo Fisher Scientific, A-11012). Cells were washed 6 times with PBST for 5 min each, and were counterstained with 300 nM of 4′,6′-diamidino-2-phenylindole (DAPI). Cells were placed on coverslips and were washed with PBS and Milli-Q water and then were mounted on glass slides with ProLong Gold mounting medium (Thermo Fisher Scientific, P36934). Cells were analyzed with a Nikon C2 confocal microscopy system (Nikon).

### Quantitative RT-PCR

Total RNA was extracted with RNeasy Plus Mini Kits (Qiagen, 74134) and reverse-transcribed with SuperScript IV (Thermo Fisher Scientific, 18090010) using random primers (Promega, C1181). Expression levels of mRNAs encoding Oct3/4, Nanog, and β-actin were analyzed by real-time PCR on a LightCycler (Roche Diagnostics) using the LightCycler FastStart DNA Master SYBR Green I kit (Roche Diagnostics, 03003230001). The amplification condition for Oct3/4 was 10 min at 95ºC for one cycle, followed by 40 cycles of 10 s at 95ºC, 5 s at 60ºC, and 10 s at 72ºC. The conditions for Nanog and β-actin were similar except that the extension step was 20 s at 72ºC for Nanog and the annealing step was 5 s at 55ºC for β-actin. Primer sequences were as follows: Oct3/4, forward: 5’-CCTGGAATCGGACCAGGCTCAGAGGTATTG-3’, reverse: 5’-ATTGTTGTCGGCTTCCTCCACCCACTTCTC-3’; Nanog, forward: 5’-CCACAGTTTGCCTAGTTCTGAGGAAGCATC -3’, reverse: 5’-TACTCCACTGGTGCTGAGCCCTTCTGAATC-3’; β-actin, forward: 5’-CAGGGTGTGATGGTGGGAATGGGTCAGAAG-3’, reverse: 5’-TACGTACATGGCTGGGGTGTTGAAGGTCTC-3’. The quantity of each transcript was measured from a standard curve, and the amounts of Oct3/4, Nanog transcript were normalized to β-actin transcript levels.

### Chromatin immunoprecipitation (ChIP) assay

The ChIP assay was carried out as described previously with some modifications[83, 84]. Briefly, cells were fixed by adding methanol-free 16% formaldehyde to the cell culture medium to a final concentration of 1% with gentle shaking at 25ºC for 10 min. After fixation of cells, 2.5 M glycine solution was added to the medium to a final concentration of 0.15 M and incubated at 25ºC for 5 min. Cells were washed twice and suspended in PBS and then were collected by scrapping into tubes. Cells were further collected by centrifugation at 300 × g at 4ºC for 5 min. The collected cells were snap-frozen in liquid nitrogen and were stored in a -80ºC deep-freezer until use. Before use, cells were defrosted on ice for 10 min. To prepare the nuclear extract, lysis buffer 1 containing 50 mM HEPES-KOH (pH 7.5), 140 mM NaCl, 1 mM EDTA, 0.25% (v/v) Triton X-100, 0.5% (v/v) NP40, 10% (v/v) glycerol, 10 mM Sodium-Butyrate, 0.1 mM PMSF, and cOmplete™ EDTA-free protease inhibitor cocktail was added to the defrosted cells and incubated on ice for 10 min. Cells were then collected by centrifugation at 800 × g at 4ºC for 5 min. Cell pellets were resuspended in lysis buffer 2 containing 50 mM HEPES-KOH (pH 7.5), 200 mM NaCl, 1 mM EDTA, 10 mM sodium butyrate, 0.1 mM PMSF and cOmplete™ EDTA-free protease inhibitor cocktail and incubated on ice for 10 min. Cells were collected again by centrifugation as described above. Finally, cells were extracted with lysis buffer 3 containing 50 mM HEPES-KOH (pH 7.5), 140 mM NaCl, 1 mM EDTA, 0.1% (w/v) sodium deoxycholate, 1% (w/v) SDS, 1% (v/v) Triton X-100, 10 mM sodium butyrate, 0.1 mM PMSF, and cOmplete™ EDTA-free protease inhibitor cocktail and were incubated on ice for 30 min. A four-times volume of dilution buffer containing 50 mM HEPES-KOH (pH 7.5), 140 mM NaCl, 1 mM EDTA, 0.1% (w/v) sodium deoxycholate, 1% (v/v) Triton X-100, 10 mM sodium butyrate, 0.1 mM PMSF, and cOmplete™ EDTA-free protease inhibitor cocktail was added to the nuclear extract to bring the final concentration of SDS to 0.2% (w/v). To prepare the nuclear extract, DNA was sonicated using Bioruptor (Diagenode) on high power under the following condition: 15 cycles of 30 s of on and 30 s of off, cooling samples on ice every 5 cycles. After the sonication step, the nuclear extract was collected by centrifugation at 20,000 × g for 10 min at 4ºC. DNA concentration was estimated by UV absorbance at 260 nm. Nuclear extracts containing an equal amount of DNA were prepared in tubes, and then an equal volume of dilution buffer was added to bring the final SDS concentration to 0.1% (w/v). For immunoprecipitation, 5 μg of anti-Histone H3K9me3 (2F3) and anti-H3K9ac (1qE5) antibodies (provided by H. Kimura) [84] and an equal amount of normal mouse IgG (Santa Cruz, sc-2025) were added to the nuclear extract and incubated overnight at 4ºC with gentle rotation. The next day, pre-equilibrated Dynabeads M-280 sheep anti-mouse IgG (Thermo Fisher Scientific, 11201D) was added to the reaction mixture and further incubated for 4 h at 4ºC with gentle rotation. Antibody-bound proteins were collected with a magnet and washed for 10 min each at 4ºC with gentle rotation in wash buffer as described below. The bound proteins were washed with low salt wash buffer containing 50 mM HEPES-KOH (pH 7.5), 140 mM NaCl, 1 mM EDTA, 0.1% (w/v) sodium deoxycholate, 1% (v/v) Triton X-100, 0.1% (w/v) SDS, 10 mM sodium butyrate, 0.1 mM PMSF, and cOmplete™ EDTA-free protease inhibitor cocktail; high salt wash buffer containing 10 mM Tris-Cl (pH 8.0), 500 mM NaCl, 1 mM EDTA, 0.1% (w/v) sodium deoxycholate, 1% (v/v) Triton X-100, 0.1% (w/v) SDS, 10 mM sodium butyrate, 0.1 mM PMSF, and cOmplete™ EDTA-free protease inhibitor cocktail; LiCl wash buffer containing 10 mM Tris-Cl (pH 8.0), 250 mM LiCl, 1 mM EDTA, 0.1% (w/v) sodium deoxycholate, 1% (v/v) Triton X-100, 0.1% (w/v) SDS, 10 mM sodium butyrate, 0.1 mM PMSF, and cOmplete™ EDTA-free protease inhibitor cocktail; and TE wash buffer containing 10 mM Tris-Cl (pH 8.0), 1 mM EDTA, 10 mM sodium butyrate, 0.1 mM PMSF, and cOmplete™ EDTA-free protease inhibitor cocktail. After a final wash with TE buffer, antibody-bound protein complexes were reverse cross-linked with elution buffer containing 10 mM Tris-Cl, pH 8.0, 300 mM NaCl, 5 mM EDTA, and 0.5% (w/v) SDS by heating at 65ºC overnight. Reverse cross-linked DNA was further treated with RNase A and Proteinase K and extracted using phenol-chloroform-isoamyl alcohol and chloroform-isoamyl alcohol as described above. Finally, the extracted DNA was precipitated with ethanol and dissolved with 10 mM Tris-Cl (pH 8.0) buffer and subjected to real-time PCR analysis as follows. A serial dilution of input DNA and antibody-bound DNA were prepared from three independent ChIP experiments and analyzed two times using a StepOne Plus real-time PCR system (Thermo Fisher Scientific). PCR cycling conditions are described below. For Oct3/4 detection, 2 min at 50ºC, 2 min at 95ºC, and 50 cycles of 15 s at 95ºC, 15 s at 53ºC, and 1 min at 72ºC. For Nanog and Sox2 detection, 2 min at 50ºC, 2 min at 95ºC and 40 cycles of 15 s at 95ºC, 15 s at 58ºC, and 1 min at 72ºC. For the U3 region of IAP detection, 2 min at 50ºC, 2 min at 95ºC, and 40 cycles of 15 s at 95ºC, 15 s at 60ºC, and 1 min at 72ºC. For the 5’ UTR region of IAP detection, 2 min at 50ºC, 2 min at 95ºC, and 40 cycles of 15 s at 95ºC, 15 s at 62ºC, and 1 min at 72ºC. For Line L1 ORF2 detection, 2 min at 50ºC, 2 min at 95ºC, and 40 cycles of 15 s at 95ºC, 15 s at 58ºC, and 1 min at 72ºC. For L1MdF detection, 2 min at 50ºC, 2 min at 95ºC, and 40 cycles of 15 s at 95ºC, 15 s at 60ºC, and 1 min at 72ºC. Primer sequences were as follows: Oct3/4, forward: 5’-ATCCGAGCAACTGGTTTGTG-3’, reverse: 5’-AAACTGAGGCGAGCGCTATC-3’; Nanog, forward: 5’-GGGTAGGGTAGGAGGCTTGA-3’, reverse: 5’-CGGCTCAAGGCGATAGATT-3’; Sox2, forward: 5’-CCTAGGAAAAGGCTGGGAAC-3’, reverse: 5’-GTGGTGTGCCATTGTTTCTG-3’; U3 region of IAP, forward: 5’-CGAGGGTGGTTCTCTACTCCAT-3’, reverse: 5’-GACGTGTCACTCCCTGATTGG-3’; 5’ UTR region of IAP, forward: 5’-CGGGTCGCGGTAATAAAGGT-3’, reverse: 5’-ACTCTCGTTCCCCAGCTGAA-3’; Line L1 ORF2, forward: 5’-TTTGGGACACAATGAAAGCA-3’, reverse: 5’-CTGCCGTCTACTCCTCTTGG-3’; L1MdF, forward: 5’-GCATCTCTGGGGTGAGCTAG-3’, reverse: 5’-AAAAGGGTGCTGCCTCAGAA-3’.

### Image analysis

Captured images of embryoid bodies and DAPI foci were binarized using the Fiji-Image J software and Photoshop CS5.1. The surface area of embryoid bodies and circularity of DAPI foci were further analyzed using the Fiji-Image J software. Circularity was calculated by 4π(area/perimeter^2). This value varies between 0 and 1. A value of 1 indicates a perfect circle.

### Statistical analysis

Statistical analyses were performed using the Pairwise t-test with Bonferroni correction. Differences were considered significant at *p*-values < 0.05.

## Acknowledgments

We thank Dr. K. Ura for assisting sucrose gradient sedimentation assay. This work was supported by MEXT KAKENHI Grant Numbers JP18K14625 (K.K.), JP18K19275 (K.H.), and JSPS KAKENHI Grant Number JP20H03174 (K.H.). This work was also supported in part by Nara Medical University Grant-in-Aid for Collaborative Research Projects (K.H.) and a research grant from the Naito Foundation (K.H.) and the Takeda Science Foundation (K.H.).

## Author contributions

K.K. and K.H. conceived and designed the project. J.Y. and K.H. generated Smarce1 mutant ES cells. J.Y. conducted expression analysis of pluripotency genes. K.K. conducted all other experiments. H.K. provided histone antibodies and assisted interpretation of the data. K.K. and K.H. wrote the manuscript with input from the other authors.

