## Supplementary_Figures for "Disruption of Smarce1, a component of the SWI/SNF chromatin remodeling complex, decreases nucleosome stability in mouse embryonic stem cells and impairs differentiation"

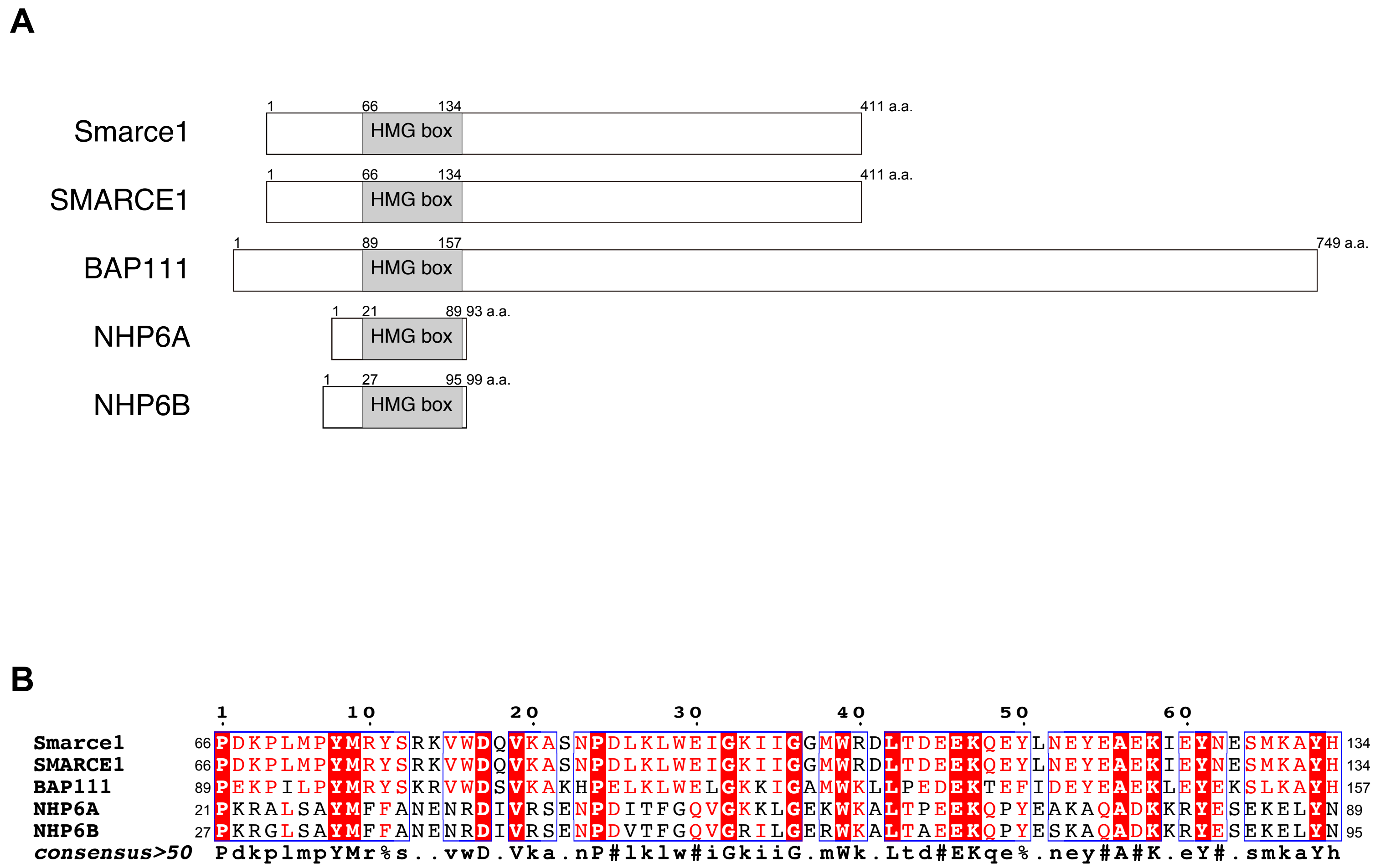


**Supplementary Figure 1. Amino acid sequence similarity between Smarce1 and other HMG box-containing proteins.**

(A) Schematic representation of HMG box-containing proteins: mouse Smarce1 (O54941), human SMARCE1 (Q969G3), Drosophila BAP111 (Q9W384), S. cerevisiae NHP6A (P11632), and S. cerevisiae NHP6B (P11633). The numbers above the rectangles represent amino acid numbers.

(B) Amino acid sequence alignment of the HMG box was performed using MultAlin (http://multalin.toulouse.inra.fr/multalin/multalin.html). White letters highlighted by a red background and red letters represent a perfect match and a conserved substitution between the HMG boxes, respectively.

**A**

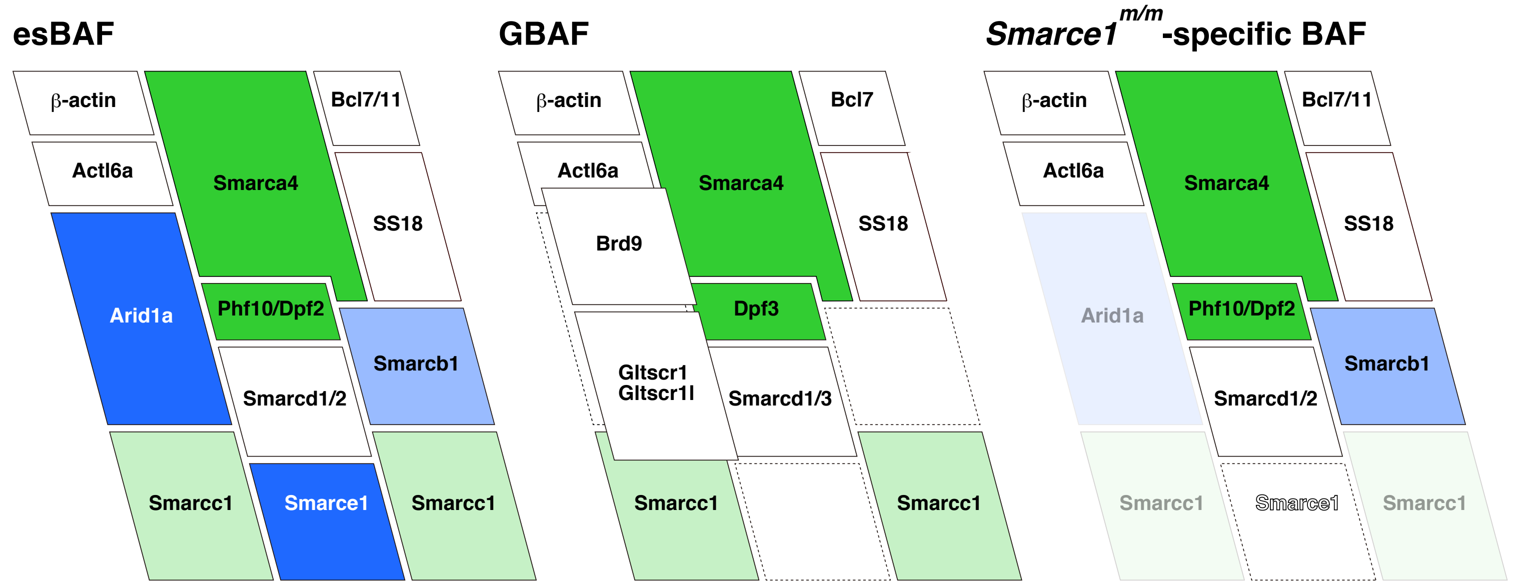


**B**

| esBAF | GBAF |
|---|---|
| Arid1a (BAF250a) | |
| Smarca4 (Brg1, BAF190a) | Smarca4 (Brg1, BAF190a) |
| Smarcc1 (BAF155) | Smarcc1 (BAF155) |
| Smarcd1(BAF60a) | Smarcd1(BAF60a) |
| Smarcd2 (BAF60b) | |
| | Smarcd3 (BAF60c) |
| Smarce1 (BAF57) | |
| Actl6a (BAF53a) | Actl6a (BAF53a) |
| Smarcb1 (BAF47) | |
| Phf10 (BAF45a) | |
| Dpf2 (BAF45d) | |
| | Dpf3 (BAF45c) |
| Bcl11a/b | |
| Bcl7a/b/c | Bcl7b/c |
| SS18 | SS18 |
| | Brd9 |
| | Gltscr1 |
| | Gltscr1l |

**Supplementary Figure 2. Component proteins of BAF complexes.**

(A) Schematic representation of component proteins of esBAF, GBAF, and *Smarce1*$^{m/m}$-specific BAF complexes. Blue and green represent experimentally validated binding capacities to DNA and histones, respectively. Light blue and green represent predicted binding capacities to DNA and histones, respectively. Unstable associations of Arid1a and Smarcc1 in *Smarce1*$^{m/m}$-specific BAF complex are illustrated with faint blue and green.

(B) The component proteins of esBAF and GBAF are listed. Blank columns indicate proteins that are not present in each complex.

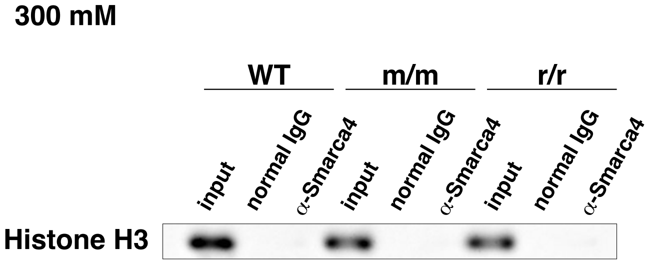


**Supplementary Figure 3. Lack of association between Smarca4 and Histone H3.** Immunoblot analysis of Smarca4-associated chromatin using anti-Histone H3 antibodies. Immunoprecipitation was conducted at 300 mM NaCl. The input represents 10% of nuclear extracts.
